# Complex relationship among vessel diameter, shear stress and blood pressure controlling vessel pruning during angiogenesis

**DOI:** 10.1101/2023.09.13.557676

**Authors:** Vivek Kumar, Prashant Kumar, Takao Hikita, Mingqian Ding, Yukinori Kametani, Masanori Nakayama, Yosuke Hasegawa

**Author notes:** Correspondence to and. These authors equally contributed. These authors equally contributed (Vivek Kumar, Prashant Kumar, Takao Hikita).

## Abstract

Blood vessel pruning during angiogenesis is the optimization process of the branching pattern to improve the transport properties of a vascular network. Recent studies show that part of endothelial cells (ECs) subjected to lower shear stress migrate toward vessels with higher shear stress in opposition to the blood flow for vessel regression. While dynamic changes of blood flow and local mechano-stress could coordinately modulate EC migration for vessel regression within the closed circulatory system, the effect of complexity of haemodynamic forces and vessel properties on vessel pruning remains elusive. Here, we reconstructed a 3-dimentsional (3D) vessel structure from 2D confocal images of the growing vessels in the mouse retina, and numerically obtained the local information of blood flow, shear stress and blood pressure in the vasculature. Moreover, we developed a predictive model for vessel pruning based on machine learning. We found that the combination of shear stress and blood pressure with vessel radius was tightly corelated to vessel pruning sites. Our results highlighted that orchestrated contribution of local haemodynamic parameters was important for the vessel pruning.

**Authors Summary:** Blood vessel networks formed by angiogenic vessel growth subsequently undergo extensive vascular remodeling process by regression of selected vascular branches. Optimization of the branching pattern in the vasculature is critical to ensure sufficient blood supply throughout the entire tissue. Recent studies have highlighted a strong relationship between the vessel remodeling and the shear stress acting on the vessel wall. However, its detailed mechanisms remain elusive due to the difficulties of estimating local haemodynamic parameters and relating them to vessel remodeling.

Here, we have numerically simulated local haemodynamic parameters within the vascular network of the postnatal day 6 (P6) mouse retinal vasculature. Then, the relationship among the local shear stress, blood pressure, and vessel radius with the vessel pruning was examined. Moreover, we developed a predictive model for the vessel pruning based on the local haemodynamic parameters by a machine learning technique. Importantly, our results indicate that the combination of shear stress and blood pressure with vessel radius is tightly correlated to vessel pruning sites.

Given the ongoing clinical approach to suppress tumor growth via blood vessel normalization, our results provide important knowledge for developing future medicine such as nanomedicine based on drug delivery systems.

## Introduction

A closed circulatory system in our body is comprised of the heart and the vasculature and is essential for development and tissue homeostasis to efficiently distribute nutrients, gases, liquids, signaling molecules and circulating cells. The blood is pumped out from the heart to the entire body through a complex hierarchical vascular network composed of arteries, capillaries and veins, causing the complex distribution of blood pressure and shear stress. Endothelial cells (ECs) form the inner lining of the vasculature and control the vascular branching pattern and other steps of vascular wall assembly(Potente & Makinen, 2017; Trimm & Red-Horse, 2022). As excessive blanching leads to uneven blood flow distribution in a tissue, optimization of the branching pattern in the vascular network after angiogenic vascular sprouting is critical to ensure sufficient blood supply throughout the entire tissue with a limited power consumption of the heart(Chen *et al*, 2012; Franco *et al*, 2015; Korn & Augustin, 2015; Korn *et al*, 2014; Lenard *et al*, 2015). Extensive rearrangement of vessel connectivity and endothelial specialization occur to ensure appropriate remodeling and quiescence of ECs. Vessel pruning leads to the selective removal of superfluous vessels by EC migration and rearrangement, leading to optimized blood flow distribution with the less pumping power.

Recent studies have highlighted a strong relationship between the vessel remodeling and the shear stress acting on the vessel wall. It has been reported that ECs tend to migrate from vessels with low shear stress to those with higher shear stress in opposition to the blood flow (Chen *et al*., 2012; Franco *et al*., 2015; Franco *et al*, 2016; Korn *et al*., 2014; Lenard *et al*., 2015; Udan *et al*, 2013). In the zebrafish midbrain and the cranial division of the internal carotid artery on eye during development, vessel pruning driven by haemodynamic parameters is observed in the developing vasculatures, leading to gradual reduction of the vascular complexity(Chen *et al*., 2012). Importantly, some of the pruning sites correspond to low shear stress regions which are predicted by numerical simulation (Bernabeu *et al*, 2018; Bernabeu *et al*, 2014; Franco *et al*., 2015; Zhou *et al*, 2021). In the mouse retina, endothelial Partitioning defective 3 (PAR-3) plays a critical role in endothelial sensitization to the wall shear stress. The loss of PAR-3 in ECs causes defected axial polarization induced by the shear stress(Hikita *et al*, 2018), resulting in enhancement of the vessel pruning. In contrast, ECs subjected to low shear stress do not always migrate toward high shear regions(Fonseca *et al*, 2020). In tumors, blood vessels are unevenly distributed with a chaotic pattern and often exhibit irregular branching(Carmeliet & Jain, 2011; Gerald *et al*, 2013). Not all open vessels are perfused continuously, and the blood flow distribution may change within a few minutes and the flow direction could be reversed even in the same vessel(Carmeliet & Jain, 2011). Dynamic changes of by blood flow should affect not only local shear stress but also other factors including blood pressure or geometry of vessels. However, the combined effect of local haemodynamic factors on vessel regression to vascular network optimization remains elusive.

Here, we have reconstructed three-dimensional (3D) vessel structures from 2D confocal images of the postnatal day 6 (P6) mouse retinal vasculature and numerically simulated blood flow to obtain local haemodynamic parameters such as wall shear stress and blood pressure within the vascular network. We further investigated the relationship among the local haemodynamic parameters including shear stress, blood pressure, and vessel radius with the vessel pruning. Moreover, we developed a predictive model for the vessel pruning based on the local haemodynamic parameters by leveraging machine learning techniques, and the established model was validated with test samples.

## Results

### Impact of vessel pruning on flow field behavior in the vascular network

Angiogenic vascular growth is important to improve tissue hypoxia. To investigate whether vessel pruning contributes to the increase of blood flow at the angiogenic front, 3D vascular structures were reconstructed from 2D confocal images using an antibody against ICAM-II, the marker for the endothelial luminal surface as previously established (Mirzapour-Shafiyi *et al*, 2021). Regressing ECs integrate into neighboring vessels and leave empty sleeves with the EC basement membrane, including collagen type IV as a remnant of the existed vascular connection(Adams & Alitalo, 2007; Baeyens *et al*, 2016; Baluk *et al*, 2003; Gerhardt *et al*, 2003; Potente *et al*, 2011). The structure containing both ICAM-II and collagen type IV positive signal were assumed as a pre-pruning structure, whereas ICAM-II positive structure is defined as a post-pruning structure (Fig. 1A). The blood is supplied from an artery at the center of the retina and is distributed to the azimuthal direction through branching capillary networks to the angiogenic front where ECs are subjected to VEGF signaling, then flows into a draining vein toward the outlet. The spatial distribution of the blood flow can be considered as a key quantity to evaluate the transport properties of the vasculature. Blood flow was numerically simulated with the pre-pruning and post-pruning blood vessel networks and the obtained blood flow intensity (magnitude of the cartesian velocity components) in each network are shown (Fig. 1B). To examine the effects of the vessel pruning on the transport properties of the vascular network, the blood flow intensity of the pre-pruning structure was subtracted from that of the post-pruning structure. Blood flow around the angiogenic front was enhanced after vessel pruning (Fig. 1C). The comparison of the pressure distributions between the pre-pruning and post-pruning vascular networks were shown in Fig. 1D. Off note that the Dirichlet boundary condition (p = 0) was employed at the outlet in both structures, so that the plotted pressure represented the relative pressure from the outlet pressure. Furthermore, the difference between the local pressures in the post- and pre-pruning structures, i.e., p_post_ – p_pre_, was plotted in Fig. 1E. The pressure was observed to be increased especially near the inlet after the vessel pruning.

**Fig 1.**
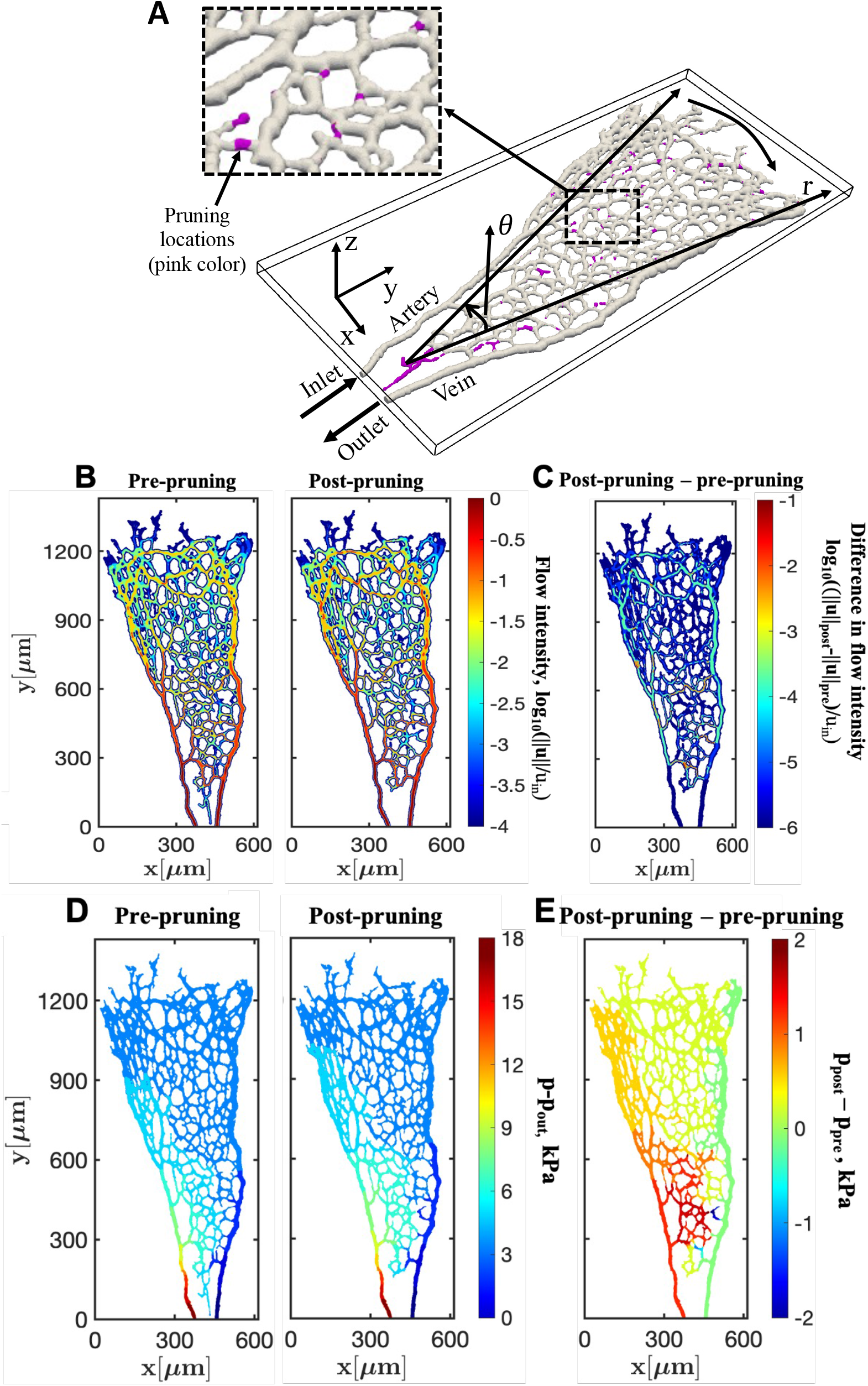
The effect of vessel pruning on haemodynamic parameters. **(A)** The 3D structure of the vascular network with pruned vessels. Gray: the structure after pruning and Pink: the pruned blood vessels. **(B)** Blood flow intensity of the pre-pruning and the post-pruning structures. Color scale represented the logarithmic normalized blood flow intensity (log_10_(∥***u***∥/*u*_*in*_)). **(C)** The difference in the flow intensities between the post- and pre-pruning structures in a logarithmic scale (log_10_((∥***u***∥_post_ - ∥***u***∥_pre_)/*u*_*in*_)). **(D)** Pressure distribution of the pre-pruning and the post-pruning structures. Color scale referred to the relative pressure to the outlet pressure, i.e., p-p_out_. **(E)** The difference between the pressures of the post- and the pre-pruning structures, i.e., p_post_ - p_pre_.

### Effects of wall shear stress on the vessel pruning

To gain further insights into the effects of vessel pruning on the transport property of the vasculature, we next analyzed wall shear stress (WSS) in several mouse retina samples (Fig. S1). The distribution of WSS was simulated as previously reported(Mirzapour-Shafiyi *et al*., 2021) and was compared with the locations of the pruned vessels (Fig. 2A). Given the fact that EC subjected to low wall shear stress migrates toward to the region subjected to high shear stress(Fonseca *et al*., 2020), we aimed to visualize relative difference of local WSS. The local average of WSS (WSS_avg_) was calculated around each point within a local small window of 91.5*μm* × 91.5*μm* × 36*μm*. Then, a dimensionless WSS index (*α*) for the local shear stress is defined as 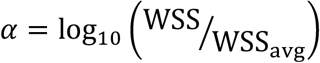. From the definition, *α* becomes positive when the local shear stress is larger than its average around that point, whereas it is negative for regions with lower shear stress. The contour of *α* in the pre-pruning vasculature was shown in Fig. 2A. The actual pruned vessel segments observed experimentally were bordered by thick black lines. Interestingly, not all of vessel branches with low WSS were regressed. Moreover, considerable amount of the vessels subjected to high WSS were also pruned (Fig. 2A). We next examined the pruning possibility (pp__*exp*_= pruned vessel wall points/total vessel wall points) obtained from the present experiment as a function of *α* (Fig. 2B). While the pruning possibility had a peak at the low shear stress of *α* = −0.9, the highest peak was observed at *α* = 0.3 where the local shear stress was relatively high. These results suggest the complex relationship between local WSS and vessel pruning.

**Fig 2.**
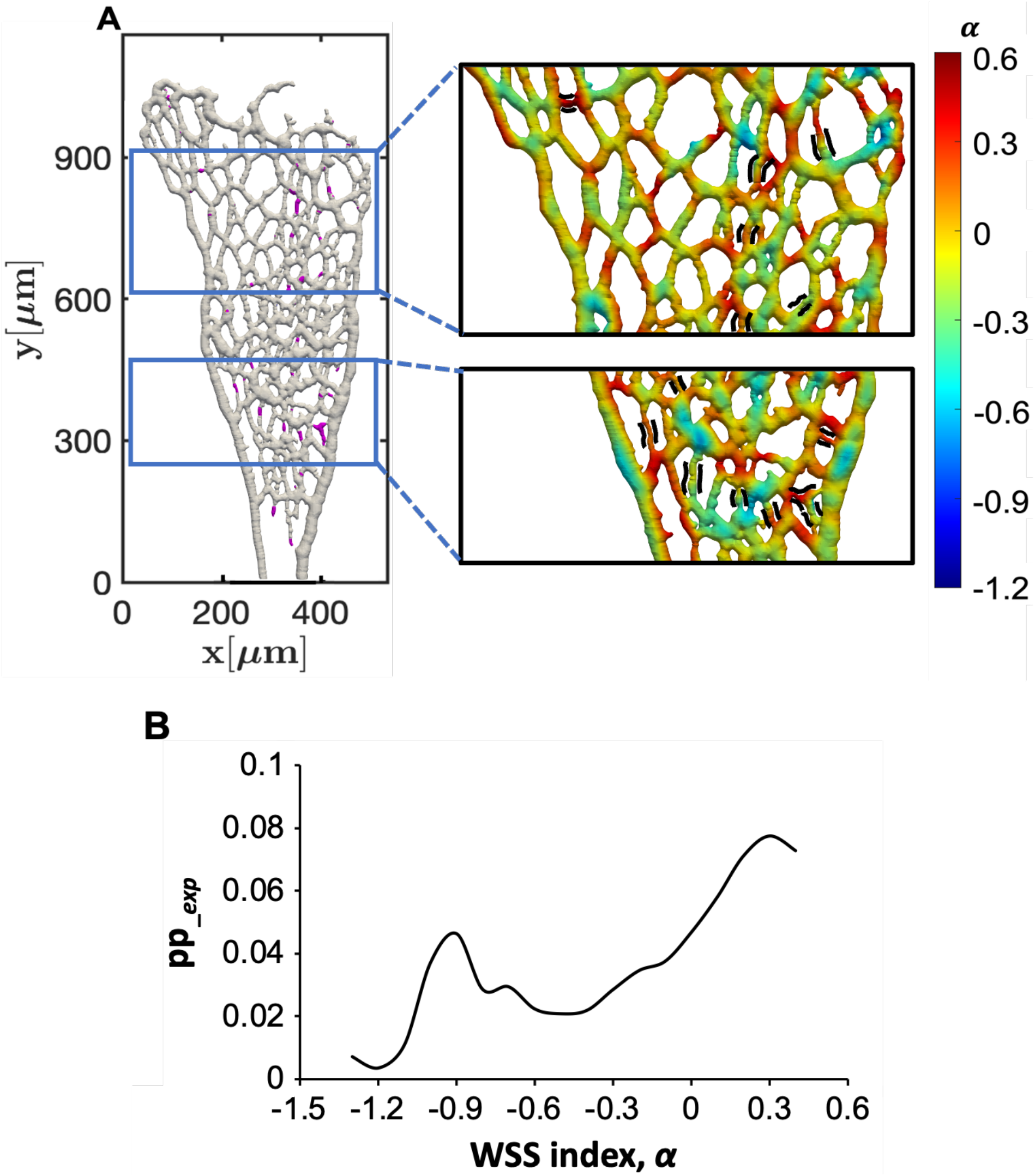
Effect of WSS on vessel pruning. **(A)** Actual pruning sites (pink) in the vascular network (gray) were shown in the left panel. Right panels showed *α* distribution in the vascular network. The color on the vessel surface represented the value of *α*. **(B)** Pruning possibility (pp__*exp*_) and *α* were plotted.

### Development of a machine learning model to predict vessel pruning sites

To further investigate the role of WSS on vessel pruning, we developed a machine learning model. Out of 7 samples (S1, S2, S3, S4, S5, S6 and S7) as shown in Fig. S1, 6 samples (S1-S6) were used for the training, whereas the remaining sample (S7) was used as a test data to validate the trained machine learning model. It was observed that pruning mostly occurred close to the inlet and outlet as well as around the angiogenic front near the vein. Considering the relative location of the pruning sites from the inlet and the outlet, we introduced a cylindrical coordinate system (*r,θ,z*) with its origin (reference point) at the first branching point from the arterial inlet as shown in Fig. 1A. The radial and azimuthal directions were denoted by *r* and *θ* respectively, while |*z*| was the third dimension normal to the (*r* -*θ*) plane. Accordingly, as the inputs of the present machine learning model, we provided the relative location from the reference point, i.e., the coordinates: *r, θ*, |*z*|, the local vessel radius (*R*) and *α* at the location of interest. Then, the model output the pixel-wise pruning possibility (Model-1). To effectively train the machine learning model, we randomly selected the same number of pixels from the unpruned regions as that of the pixels of the pruned ones (Fig. 3A). During the training, we repeated randomly selecting the data points, i.e., the pixels from the unpruned vessels, every 10 epochs, while all the points of the pruned vessels were used for the training. As a result, our model predicted 9 pruned sites among 14 actual pruned vessels in the test sample S7. In addition, Model-1 incorrectly identified 6 pruning sites. The predicted pruning sites (pixels) were determined if the pruning probability output from the model was greater than a critical value determined by maximizing the Intersection over Union (IoU) between the prediction and the ground truth. Specifically, the critical probability of 0.79 was used (Fig. 3B). A vessel is defined as pruned if pixels identified as pruned completely cover the perimeter of the vessel. For more detailed procedures to determine the pruned vessels, see Materials and Methods.

**Fig 3.**
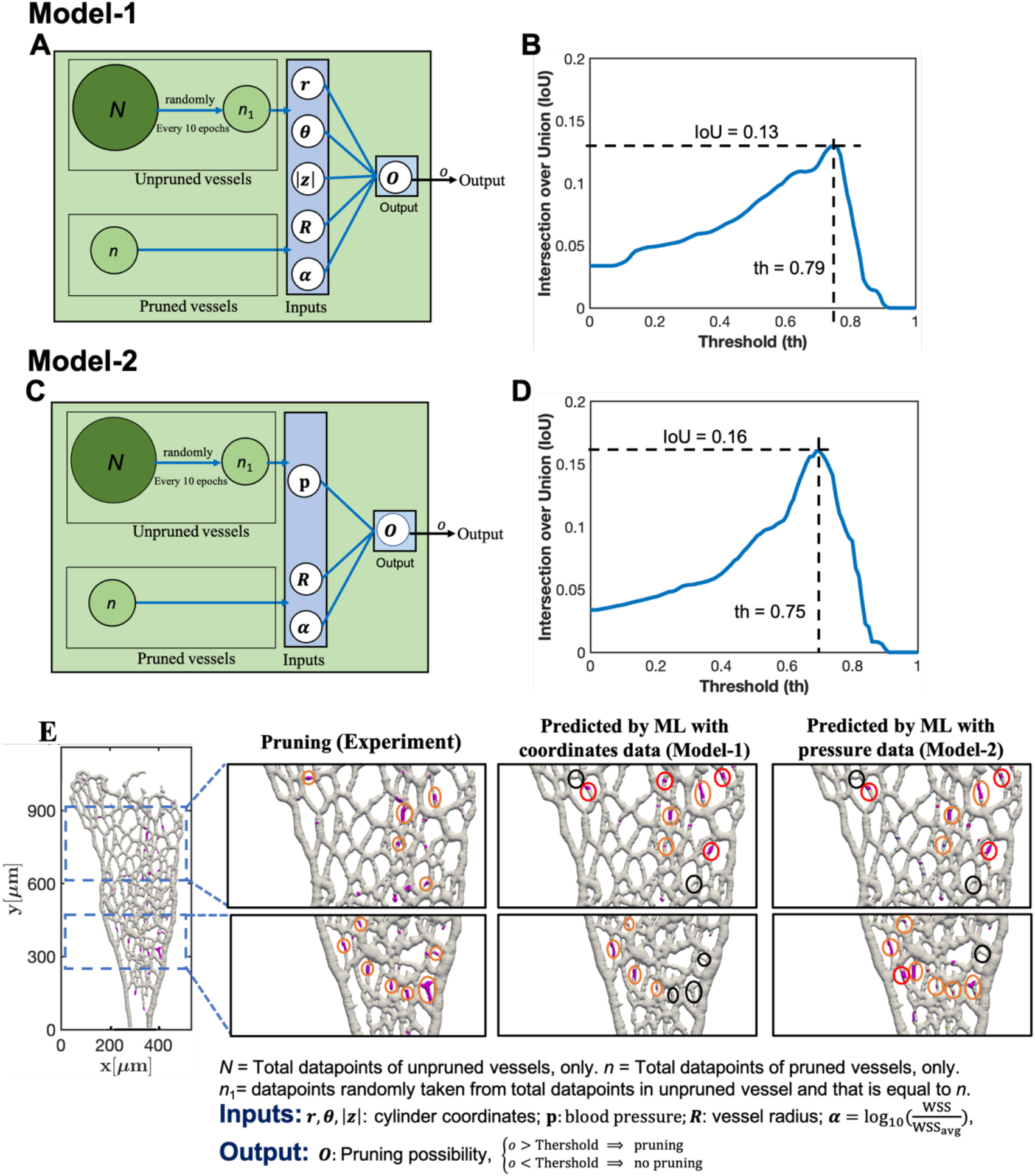
Vessel pruning prediction by machine learning model. **(A)** Schematic of the machine learning model (Model-1) in which the relative location from the inlet (*r, θ*, |*z*|), the vessel radius (*R*) and the *α* were input to the network, while the pruning possibility was output. *N* was the total number of pixels corresponding to unpruned vessels, while *n* was the total number of pixels corresponding to the pruned vessels. *n*_*1*_ was the number of pixels randomly sampled from *N* unpruned pixels every 10 epochs. **(B)** IoU as a function of the threshold value (th) of the pruning possibility output from Model-1. **(C)** Schematic of machine learning model (Model-2) in which blood pressure, vessel radius and WSS index were used as the input, and the pruning possibility was output. *N, n*_*1*_ and *n* were defined as same as in Model-1. **(D)** IoU as a function of the threshold (th) used in Model-2. (**E)** Comparison of vessel pruning predicated by machine learning models (Model-1 and Model-2) with the actual pruning sites. ML in the figure means ‘machine learning’. Orange, black and red circles illustrated the successfully predicted pruning sites, unpredicted pruning sites and extra predicted sites, respectively.

### Local blood pressure information improves the success rate of the pruning site prediction

To investigate whether the vessel pruning can be predicted solely by local haemodynamic factors, we considered using the blood pressure instead of the coordinate information. We found that the blood pressure gradually decreased from the inlet to the outlet along the flow direction (Fig. 1D) suggesting that the magnitude was implicitly related to the distance from the inlet. To incorporate this information of the local blood pressure, the machine learning model was modified. The coordinate information was removed from the input and the local pressure information was applied (Model-2, Fig. 3C). From actual 14 pruning locations, Model-2 could successfully predict 11 locations, while Model-1 only predicted 9 locations. Model-2 also predicted less extra pruning locations, i.e., 4 locations, whereas 6 locations were incorrectly predicted by Model-1. Consequently, the success rate for predicting the pruning sites was increased from 64% to 78.5% and IoU was increased from 0.13 to 0.16 (Fig. 3B, D). Most unpredicted pruning sites in Model-1 were located at the vascular plexus around the center of retina, where *α* showed relatively high values (Fig. 3E). In contrast, most of incorrectly predicted pruning sites existed around the angiogenic front in both models.

### The combination of *α*, local blood pressure and vessel radius improved pruning prediction accuracy

To compare actual pruning possibility between prediction by Model-2 and pruned vessels observed in the mouse retina (pp__*exp*_) directly, the output pruning probability of Model-2 (pp__*ML*_) was also rescaled based on the number of the total vessel wall points in each sample. The input parameters in Model-2 were WSS index *α*^*^, local blood pressure p^*^and local vessel radius R^*^. All these quantities were rescaled so that the range of the change for each variable was −1.3 ≤ *α*^*^ ≤ 0.3, 0 ≤ p^*^ ≤ 1.0, and 0 ≤ R^*^ ≤ 1.0, respectively, which was commonly used for efficient training of deep neural networks(Huang *et al*, 2023; Kissas *et al*, 2020). As mentioned in the previous section, we selected the same number of unpruned vessel points as that of pruned vessel for network training, and distribution of all the pixels consisting of the vessel walls for all the six training samples in the three-dimensional space defined by the three local input variables, i.e., WSS index *α*^*^, blood pressure p^*^ and vessel radius R^*^ (Fig. 4A). The blue and red circles corresponded to unpruned and pruned pixels, respectively. We also plotted the iso-surfaces of the pruning possibilities pp__*ML*_ = 0.05, 0.06, 0.07 and 0.08 estimated from the trained network of Model-2 with different colors of red, green, magenta, and cyan, respectively. It can be confirmed that the trained network captures the general trend of the training samples quite well, suggesting the training has been successfully conducted. To validate the generality of the trained network, the same plot has been provided for the test sample. The training network predicted the pruned vessel for the unseen data well (Fig. 4B)

**Fig 4.**
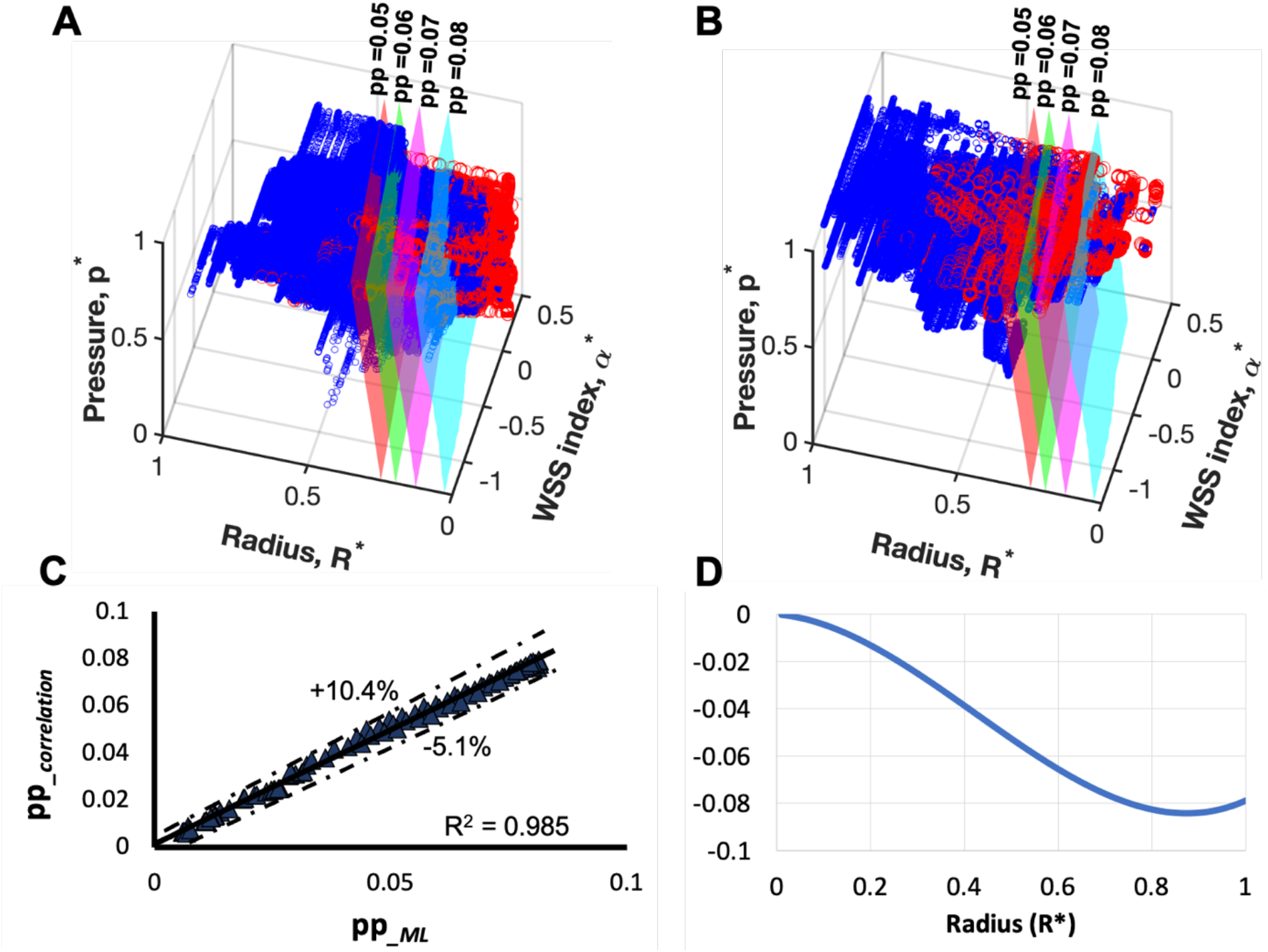
Evaluation of the model by the machine learning model with blood pressure. **(A and B)** Visualization of dependency of pruning possibility on pressure, vessel radius and WSS index. The blue and red scattered circles showed unpruned and pruned pixels from experimental data **(A)** and prediction of the pretrained network **(B)**. The iso-surfaces illustrated the pruning possibility (pp) = 0.05(red), 0.06(green), 0.07(magenta) and 0.08(cyan) that predicted from pre-trained network. p* and R* indicated the scaled pressure and radius. *α*\* referred to the scaled WSS index. **(C)** The variation of predicted pruning possibility by correlation (pp__*correlation*_) with the pruning possibility by machine learning algorithm(pp__*ML*_). **(D)** The contribution of the vessel radius to the pruning probability, i.e., 0.231R^*3^ − 0.2946R^*2^ − 0.016R^*^ appearing in Eq. (1), as a function of the dimensionless vessel radius R^*^.

Furthermore, the iso-surfaces of the pruning possibility obtained by the trained network indicated that the pruning possibility increased with increasing the WSS and the blood pressure, and decreasing the vessel radius. Specifically, the output, i.e., the pruning probability of Model-2, can be correlated with the following simple formula with the R-square value of 0.985:

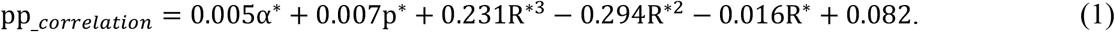

The output of Model-2, pp__*ML*_, and the proposed correlation, pp__*correlation*_ obtained from Eq. (1) were compared in Fig. 4C. The proposed correlation provided satisfactory estimate of the output of Model-2 within the error between −5.1% to +10.4%. From Eq. (1), we can conclude that the higher WSS increased the pruning possibility, since its proportional constant was 0.005, and positive. Similarly, a larger local pressure promoted more vessel pruning, and this trend was even stronger than that of WSS, as the proportional constant for the local pressure was larger. The contribution of the vessel diameter to the pruning probability, i.e., 0.231R^*3^ − 0.294R^*2^ − 0.016R^*^, was plotted as a function of the dimensionless radius (Fig. 4D). These results suggest that the pruning probability decreases with the increase in the vessel radius.

Next, we investigated the contribution of each haemodynamic factor on the pruning possibility based on Eq. (1). The pruning possibilities obtained by the experiment and Model-2 were plotted as a function of WSS index (*α*\*) together with the contribution from each haemodynamic factor on the right-hand-side of Eq. (1) (Fig. 5A). According to Eq. (1), *α*\* increased the pruning possibility linearly (Fig.5A, the blue solid line with triangles), whereas the experimental result showed a non-monotonic trend with respect to *α*\* (Fig.5A, the black line). This discrepancy can mostly be explained by the contribution from R* which showed non-monotonic dependency on *α*\*(Fig.5, the blue line with diamonds). As a result, by summing up all the contributions, the non-monotonic dependence of the experimental data (pp__*exp*_) can be predicted by the correlation (pp*_*_*correlation*_) (Fig.5A, the blue line). To gain further insight into the contribution of R* and *α*\*on pp__*correlation*_, their joint probability density function (PDF) was plotted and the most probable value of R* at each *α*\* was shown (Fig. 5B). As smaller R* was seen at the vessels with both lower and higher *α*\*, pp__*correlation*_ resulted in non-monotonic dependency on *α*\* (Fig. 5A).

**Fig 5.**
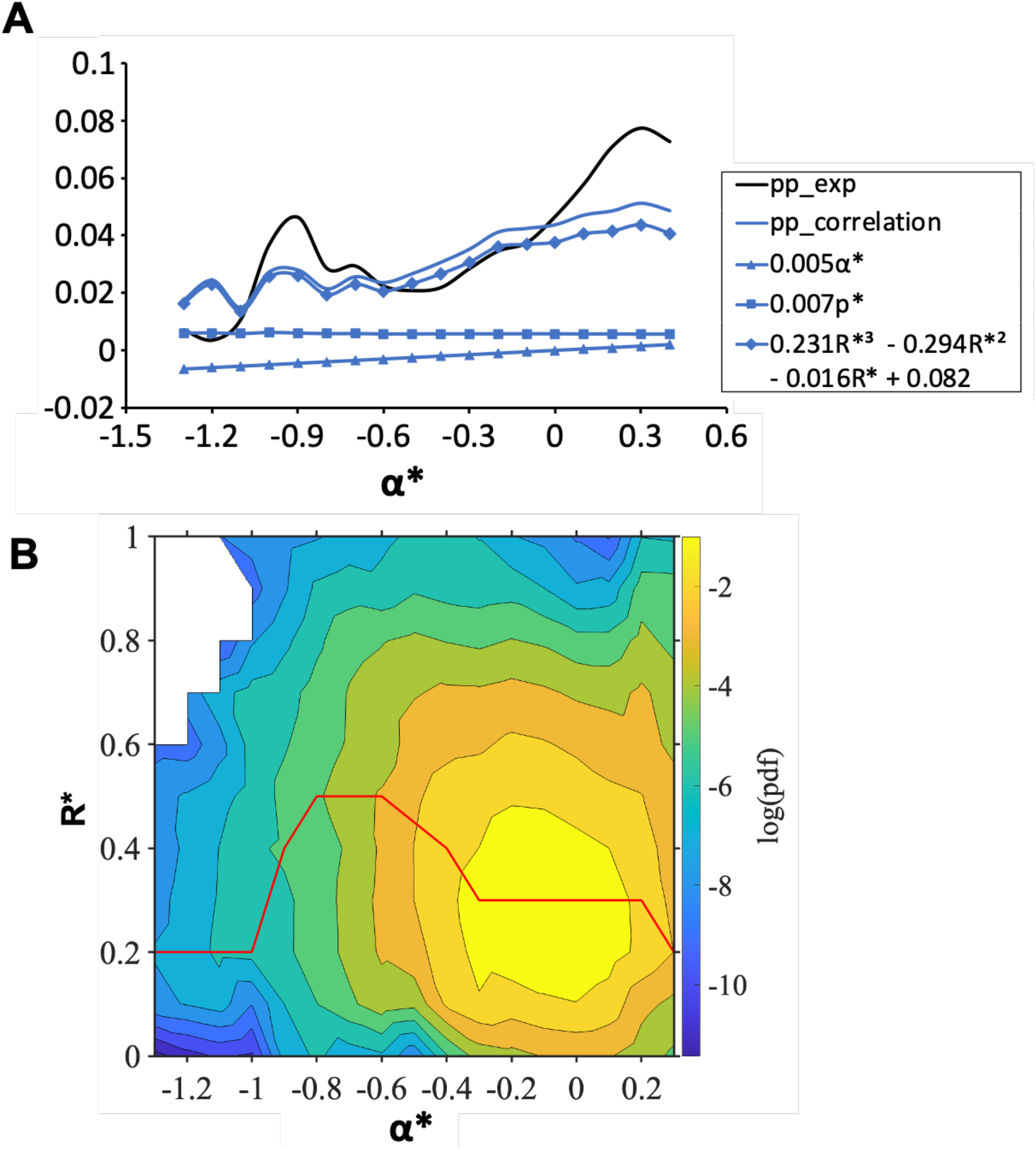
Contribution of haemodynamic factors on vessel pruning sites. **(A)** Comparison of pruning possibility by correlation (pp__*correlation*_) and experimental pruning possibility (pp__*exp*_). All contributed factors to pp__*correlation*_ were plotted against *α*^*^ with blue lines with different solid symbols. **(B)** Joint probability density function (PDF) of the normalized WSS index *α*^*^ and vessel radius R^*^. The red line shows the maximum location of the joint PDF at each *α*^*^.

## Discussion

Previous studies demonstrated the importance of WSS on vessel pruning (Korn & Augustin, 2015). During angiogenesis, vessel regression is induced by polarized endothelial cell migration. ECs subjected to low WSS are known to migrate toward the high WSS region for vessel pruning(Franco *et al*., 2015). For polarized EC migration, dynamic changes of microtubules are observed with continuous blood flow (Chiu & Chien, 2011; Hahn & Schwartz, 2009). EC shape is elongated along the axis of blood flow and migration. Microtubules are stabilized toward the direction of cell migration and the microtubule organization centers (MTOCs) and the Golgi apparatus are aligned in front of the nuclei toward the direction of migration. The loss of cell polarity factor, PAR-3 in ECs attenuates microtubules stabilization and polarization along the axis of blood flow(Hikita *et al*., 2018). As a result, vessel regression is compromised suggesting importance of WSS on determining vessel pruning sites in the vascular network(Nakayama *et al*, 2013). However, we showed that only a limited fraction of ECs subjected to low WSS underwent vessel pruning. Moreover, the vessels even exposed to high WSS were often regressed (Fig. 2B). To gain further insight into the role of mechano-stress on vessel pruning, we have developed two machine learning models to predict the pruning possibility of the growing vasculature in the mouse retina. In addition to vessel diameter and local WSS, Model-2 configurated with the local blood pressure information, was compared to Model-1 which estimated the pruning probability based on the relative location of the point of interest from the inlet. As a result, Model-2 incorporated the local blood pressure information showed improved efficiency for the prediction, suggesting the importance of blood pressure during vessel pruning.

Consistently, recent studies have shown that blood flow promotes lumen formation by inducing the formation of inverse membrane blebs during angiogenesis(Gebala *et al*, 2016). Moreover, intraluminal pressure (IP) driven by blood flow is shown to be critical for wound angiogenesis in zebrafish and in vitro cell culture(Yuge *et al*, 2022). During wound angiogenesis, blood flow-driven IP loading inhibits elongation of injured blood vessels located at sites upstream from blood flow, while downstream injured vessels actively elongate. IP induces actin cytoskeletal reorganization in ECs under Rho family small GTPase, Cdc42(Yuge *et al*., 2022). Since ECs are exposed to not only shear stress but also blood pressure, the orchestration of microtubules stabilization and actin cytoskeletal reorganization would be important for vessel pruning. Signaling crosstalk among Rho family small GTPases such as RhoA, Rac1 and Cdc42 is tightly controlled downstream of EC-to-EC junctions, integrins or primary cilia, modulating microtubule stabilization and actin cytoskeletal rearrangement (Jaffe & Hall, 2005; Nakayama *et al*, 2008; Ridley, 2015; Ridley *et al*, 2003). Interestingly, cell polarity factors, such as PAR-3, aPKC or PAR-6 are known to mediate signaling crosstalk among RhoA, Rac1 and Cdc42(Lin *et al*, 2000; Nakayama *et al*., 2008; Nishimura *et al*, 2005; Zhang & Macara, 2008).

Next, we investigated the contribution of each haemodynamic factor on vessel pruning based on the machine learning model incorporated with blood pressure information. The pruning possibility did not show monotonic correlation with WSS and have two peaks at both lower and higher WSS (Fig. 2B and 5A). According to Eq. (1), the contribution of vessel diameters on pruning was more than that of WSS, which explains the non-monotonic dependency of the pruning possibility on WSS (Fig. 5). Vessels with small diameters with low WSS generally have low flow rates, and might be considered to be pruned since they hardly affect the blood flow distribution in the entire vascular network. In contrast, vessels with higher shear stress carry more blood flow, and pruning of such vessels should have a greater impact on the blood flow distribution and supply more blood to the angiogenic front. Eq. (1) showed another peak for pruning possibility at the vessels subjected to very low WSS, which was not seen in the experimental data (Fig. 5). ECs at the angiogenic front, such as tip cells, are stimulated with VEGF for cell migration and seems to receive limited effect of blood flow. Further analysis will be warranted.

## Materials and methods

### Ethics statement

All the experimental procedures involving animals were conducted in accordance with the local animal ethics committees and authorities (Regierungspräsidium Darmstadt, B2/1073) and institutional rules and regulations.

### Retina staining

For the experiment, 6 days postnatal (P6) mice were utilized. At this desired stage of development, mouse eyes were collected and fixed for 5 hours in ice-cold 2% paraformaldehyde (PFA) in PBS for 5 hours at 4°C. After that, the retina samples were dissected in PBS. Blocking/permeabilisation was applied using Blocking Buffer (BB), consisting of 1% FBS (Gibco), 3% BSA (Sigma), 0.5% triton X100 (Sigma), 0.01% Na deoxycholate (Sigma), 0.02% Na zide (Sigma) in PBS of pH = 7.4 for 2-4 hours at 4°C on a rocking platform. Then they were incubated with primary antibodies (anti-ICAM-II (BD Pharmingen, 553326, 1:100), anti-Collagen-Type IV (Collagen-IV) (Bio-RAD, 2150-1470, 1:400) and anti-VEGF164 (R&D Systems, AF-493-NA, 1:100) in 1:1 BB/PBS), overnight at 4°C on rocking platform. Retinas were then washed four times for 30 min in PBS/ 0.2% TritonX-100 (PBT) at RT and incubated with Alexa Fluor conjugated secondary antibodies (Invitrogen, 1:500) in 1:1 BB/PBS for 2hr at RT. After another four times of washing with PBT, retinas were radially cut into four lobes and flat-mounted onto slides using Fluoromount-G mounting medium (Southern Biotech, 0100-01).

### Image processing and numerical simulation

To perform the three-dimensional simulations, the confocal images were converted to three-dimensional vascular network. The images were firstly transferred to binary images in which black and white regions represent tissue and blood vessel, respectively. In the next step, the shortest distance from the centerline to the vessel wall was calculated and then a 3D sphere with the diameter of the shortest distance was constructed on the point of the centerline. The envelope of 3D spheres at all the centerline points was used to reconstruct 3D vascular structure, and it was used as the computational domain for the present simulation.

A signed distance function, commonly known as level-set function, was applied to represent a complex three-dimensional vascular structure and was integrated into an in-house CFD solver, which had been successfully applied to blood flow simulations for mouse retinas in a previous study (Mirzapour-Shafiyi *et al*., 2021). The flow was assumed to be incompressible, Newtonian and steady. The Reynolds number (*Re*) was fixed and is equal to 0.1, based on inlet bulk mean velocity(*U*_*in*_),diameter of inlet artery (*D*_*in*_) and kinematic viscosity (*v*) reported previously (Brown *et al*, 2005; Windberger *et al*, 2003). The blood flow inside the complex three-dimensional vascular network was solved using a volume penalization method, where the effects of the complex geometry are expressed by an artificial body force acting around the interface between the fluid and solid regions. Uniform grids were generated in the whole computational domain with grid spacing (Δ_*x*_, Δ_*y*_, Δ_*z*_) ≈ (1.5 *μm*, 1.5 *μm*, 1.5 *μm*).

### Machine learning

In developing the current machine learning models (Model-1 and Model-2), six samples (S1-S6) were used for training, and the remaining sample (S7) was used for testing out of the total seven samples. In Model-1, the location information (cylindrical coordinates, *r, θ*, |*z*|), the WSS index (*α*) and vessel radius (R) were used as the input data. Meanwhile, in Model-2, the location information was replaced by the local pressure information, so that all the inputs were local haemodynamic and geometric parameters. In both the models, the same network was used to predict the pruning possibility at each pixel comprising the vessel wall. Although we used only six samples for training, each sample had around 1 million pixels making the vessel wall, so that the number of training points is around 6.7 million. It turned out that the number of unpruned points was about 20 times larger than the pruned points. To prevent the machine learning models from outputting a trivial solution of a zero probability for all the points, a portion of the unpruned points was randomly selected to make sure that the number of pruned and unpruned points in the training data are the same. The datapoints in unpruned vessels were reselected every 10 epochs to use all the pixels for training. The batch size was set to be 60000, which resulted in the best prediction among the other values we had tested.

As for the network structure, we first employed a feedforward neural network with 4 hidden layers and 20 nodes per layer. Leaky ReLU was used as an activation function for all the hidden layers and the sigmoid activation function for the last layer. By using the sigmoid activation function in the last layer, we were effectively constrained the network output between 0 and 1 to represent the pruning possibility. The loss function was set to be the binary cross-entropy between the grand truth and the model prediction. We had found that the resultant prediction accuracy was marginal. Then, we also tried a further simple network without hidden layers. In this case, the output pruning possibility was expressed as follows:

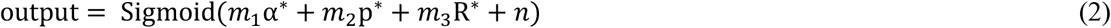

Here, *m*_1_, *m*_2_, *m*_3_ and *n* are the network parameters to be optimized by training. The above neural network provided a better prediction that those from more complicated networks. This might be attributed to the lack of the training data to train larger networks, and there is a possibility that the prediction accuracy could be further improved once more training data become available.

For the validation of the trained machine learning models, the network output which represents the pruning possibility and ranges from zero to one, has to be assessed by experimental data. We introduced the threshold value (th) for the pruning possibility and assumed that pruning occurred when the predicted possibility is larger than the value. The threshold values were determined as th = 0.79 and 0.75 for Model-1 and Model-2, respectively, to maximize Intersection of Unit (IoU) of the pixels corresponding to the pruned vessels in the experiment and the model prediction (Figs. 3(B) and (D)). Once pruned pixels with pruning possibilities higher than the threshold value were identified, they were used to predict pruned vessels. Specifically, we defined a vessel as pruned if both ends of the vessel were connected to adjacent vessels (none of the end points are open) in the pre-pruning structure, and the perimeter of the vessel was completely covered by the pruned pixels. The vessels predicted to be pruned by the present machine learning models were shown in Fig. 3 (E).

## Supporting information

**Fig S1**. 3D structures of the vascular network with pruned vessels used for training (sample no. S1-S7). Pruning locations were shown in pink color.

**Fig S2**. Validation of machine learning models. **(A)** Comparison of vessel pruning predicated by machine learning models (Model-1 and Model-2) with experimentally identified pruning sites (experiment) from sample S1. Orange, black and red circles illustrate the predicted pruning sites, unpredicted pruning sites and extra predicted sites, respectively. **(B)** Threshold values for the pruning possibility in Model-1 and Model-2.

## Acknowledgements

The present study was supported by Grants-in-Aid for Scientific Research-Fund for the promotion of Joint International Research (Fostering Joint International Research (B) and JSPS Kakenhi (JP22K08125). The sponsor had no role in the design and conduct of the study; collection, management, analysis, and interpretation of the data; preparation, review, and approval of the manuscript; or in the decision to submit the manuscript for publication. The funders had no role in study design, data collection and analysis, decision to publish, or preparation of the manuscript.The authors gratefully acknowledge former graduate students, Mr. Fumiki Mochida, Mr. Chang Cui and Mr. Mingqian Ding, at Institute of Industrial Science, The University of Tokyo for their contributions to developing the software used in this study.

